# Topological Features of Electroencephalography are Reference-Invariant

**DOI:** 10.1101/2020.09.25.311829

**Authors:** Jacob Billings, Ruxandra Tivadar, Micah M. Murray, Benedetta Franceschiello, Giovanni Petri

## Abstract

Electroencephalography (EEG) is among the most widely diffused, inexpensive, and applied neuroimaging techniques. Nonetheless, EEG requires measurements against a reference site(s), which is typically chosen by the experimenter, and specific pre-processing steps precede analysis. It is therefore valuable to obtain quantities that are reference-independent and minimally affected by pre-processing choices. Here, we show that the topological structure of embedding spaces, constructed either from multi-channel EEG timeseries or from their temporal structure, are subject-specific and robust to re-referencing and pre-processing pipelines. By contrast, the shape of correlation spaces, that is, discrete spaces where each point represents an electrode and the distance between them that is in turn related to the correlation between the respective timeseries, were neither significantly subject-specific nor robust to changes of reference. Our results suggest that the shape of spaces describing the observed configurations of EEG signals holds information about the individual specificity of the underlying individual’s brain dynamics, and that temporal correlations constrain to a large degree the set of possible dynamics. In turn, these encode the differences between subjects’ space of resting state EEG signals. Finally, our results and proposed methodology provide tools to explore the individual topographical landscapes and how they are explored dynamically. We propose therefore to augment conventional topographic analyses with an additional – topological – level of analysis, and to consider them jointly. More generally, these results provide a roadmap for the incorporation of topological analyses within EEG pipelines.

## 1 Introduction

Electroencephalography (EEG) is a non-invasive neuroimaging technique measuring the electrical activity of the brain at the scalp (Biasiucci et al., 2019). EEG has several practical strengths as a neuroimaging tool (Michel Christoph M., 2009; Michel and Murray, 2012; Michel et al., 2004; Murray et al., 2008): it is temporally precise, cost-effective, easy to use, portable and combinable with other techniques, such as MRI and PET. Indeed, these strengths made EEG a primary tool for studying brain activity both from the clinical and research standpoints (Lepage et al., 2014). EEG primarily measures postsynaptic potentials of pyramidal neurons: the neurotransmitter release generated by excitatory or inhibitory action potentials results in local currents at the apical dendrites of the post-synaptic neuron, that in turn lead to current sources and sinks in the extracellular space. In biophysical terms, voltages refer to the exertion needed to move charge from one site to another. More practically, this means that voltage is the difference between a “chosen” electrode and a “reference” electrode (Tivadar et al., 2019; Biasiucci et al., 2019). EEG is the measurement of this voltage as it varies in time, and thus results in “time series” across different sites on the scalp. Two issues arise from the biophysical underpinnings of the EEG signal. First, the brain signal recorded at scalp level is given by the synchronous activity of multiple neurons. Therefore, a given electrode not only captures brain activity from directly beneath it, but to a certain extent from the entire brain. Second, measurements of voltage are referential, meaning that EEG time series (including event-related potentials (ERPs) at a given electrode or scalp site will change when the reference changes, as there is no electrically neutral spot on the scalp (or body surface). This has led to a long-standing debate in the EEG community, discussing which of the references is more informative for the analyses (Chella et al., 2016; Yao et al., 2019; Hu et al., 2019).

This issue concerns spontaneous data as well as pre-processed and post-processed averages, and functional connectivity data. Thus, referencing affects both temporal and spatial aspects of the recorded potentials (Chella et al., 2016). With regards to spatial aspects, the values of the electric field at the scalp will change when the reference changes, as a different baseline value (i.e. the voltage of the reference electrode) is being compared to every other electrode.

In terms of temporal aspects, a non-neutral reference introduces time-varying activity into the recordings of all electrodes, meaning that both the temporal waveforms, as well as their spectral properties suffer from distortions (Chella et al., 2016). Therefore, the variance around a mean voltage value (e.g., spectral power, amplitude, etc.), as well as other derived and associated measures, including results of statistical contrasts, will change when the reference changes. These facts have generally led to –and to some extent continue to result in– misinterpretation and misuse of EEG data, despite good quality in experimental design (Michel and Murray, 2012; Biasiucci et al., 2019; Tivadar et al., 2019).

To solve the reference issue, many in the EEG community have turned towards the characterization and analysis of properties of the electrical field at the scalp, such as topographical maps and spatial pattern analysis methods, as well as source localisation techniques (MWong, 2012; Michel and Murray, 2012; Michel et al., 2004; Grave de Peralta Menendez et al., 2000; Lehmann and Michel, 2011; Michel et al., 2001; Tenke and Kayser, 2005; Marinazzo et al., 2019). Treating the data from the entire electrode montage as a multivariate vector has several advantages over waveform-based analysis of voltage. First, the shape of the electrical field at the scalp will not change with a changing reference (cf. Fig. 3 in (Tivadar et al., 2019)). Second, multivariate analyses also profit from the added information of high-density recordings. They can disentangle effects of strength from effects due to changes in sources’ configuration or signal latency (Murray et al., 2008). Third, topographic information has direct neurophysiologic interpretability (Michel and Murray, 2012), as biophysical laws dictate that differences in topography indicate changes in the configuration of active cerebral sources (Vaughan, 1982; Lehmann, 1987). Fourth, EEG mapping is the precursor for EEG source imaging (Michel and Murray, 2012). Nevertheless, traditional waveform-based analyses still predominate in the EEG community (Luck, 2014).

Here, we propose a new method of description of the EEG signal, which is robust across different pre-processing and reference choices. We first build representations of EEG data based on different type of signal embeddings. We then assess their robustness and discriminatory power, e.g. between different subjects and tasks, using recent topological data analysis tools. These tools have been shown to be useful in the analysis of neurophysiological data (Petri et al., 2014, 2013; Ibáñez-Marcelo et al., 2019a; Bassett and Sporns, 2017; Giusti et al., 2016; Varley et al., 2020) because they are built to detect properties of datasets, e.g. point clouds or weighted networks, that are invariant under homeomorphic transformations, which include, deformations, rotations, contractions and any other continuous transformation. The rationale for this ability is that they capture and quantify topology, that is, the shape of spaces in arbitrary dimensions, including discrete spaces obtained from signals, via their topological features, e.g. connected component, 1-dimensional holes, three-dimensional cavities, and the higher-dimensional analogues.

Because of this capacity, topological observables describe information qualitatively robust to noise, deformation and continuous transformations Zomorodian and Carlsson (2005); Ghrist (2008). In this way, topological descriptions of EEG data have been shown to find meaningful simplifications of high-dimensional data, by extracting low-dimensional summaries of the dataset’s shape (Giusti et al., 2015), to capture meso-scale patterns of disconnectivity (Petri et al., 2014; Lee et al., 2012) and to explicitly encode interactions among many elements (i.e. nodes in a network, regions of the brain, etc.) (Iacopini et al., 2019).

We find that embeddings constructed from multi-channel EEG timeseries, and from their temporal structure, are specific to subjects and robust to re-referencing and pre-processing pipelines. At difference, spaces obtained from spatial correlations among electrodes space, analogous to traditional functional connectivity Sporns (2013); Hutchison et al. (2013); Betzel et al. (2014), were characteristic of subjects and lacked robustness to changes of reference. Our results highlight the necessity and propose tools to explore the individual topographical landscapes and how they are explored dynamically.

## 2 Methods

### 2.1 Overview of analysis pipeline

In the following sections, we summarise the pipeline we adopt to analyse EEG data and compare the results across multiple references and pre-processing choices. We start from raw EEG signals recorded on the scalp in a cohort of 21 subjects (Fig. 1a, see section 2.2). The recorded signals are then prepared using three different pre-processing pipelines (Fig 1b, see Section 2.3). For each pipeline, we also consider different EEG references, which can result in signals with different waveforms and different relations to one another (Fig. 1c): for illustration, we show three EEG signal snippets for various references. For all pre-processing pipelines, subjects and references we compute three different representations: functional connectivity (Sec. 2.4), the Takens embedding (Sec. 2.5) and the direct temporal embedding (Sec. 2.6) (Fig. 1d). For all these representations we extract summaries of their low-dimensional topological properties (Sec. 2.7) and use them to compute distances between them (Fig. 1e-f). Finally, we compare the results obtained from the previous analysis with a synthetic benchmark obtained by temporally reshuffling the EEG data (Sec. 2.8). The following sections provide details on the analysis steps described above.

**Fig. 1.**
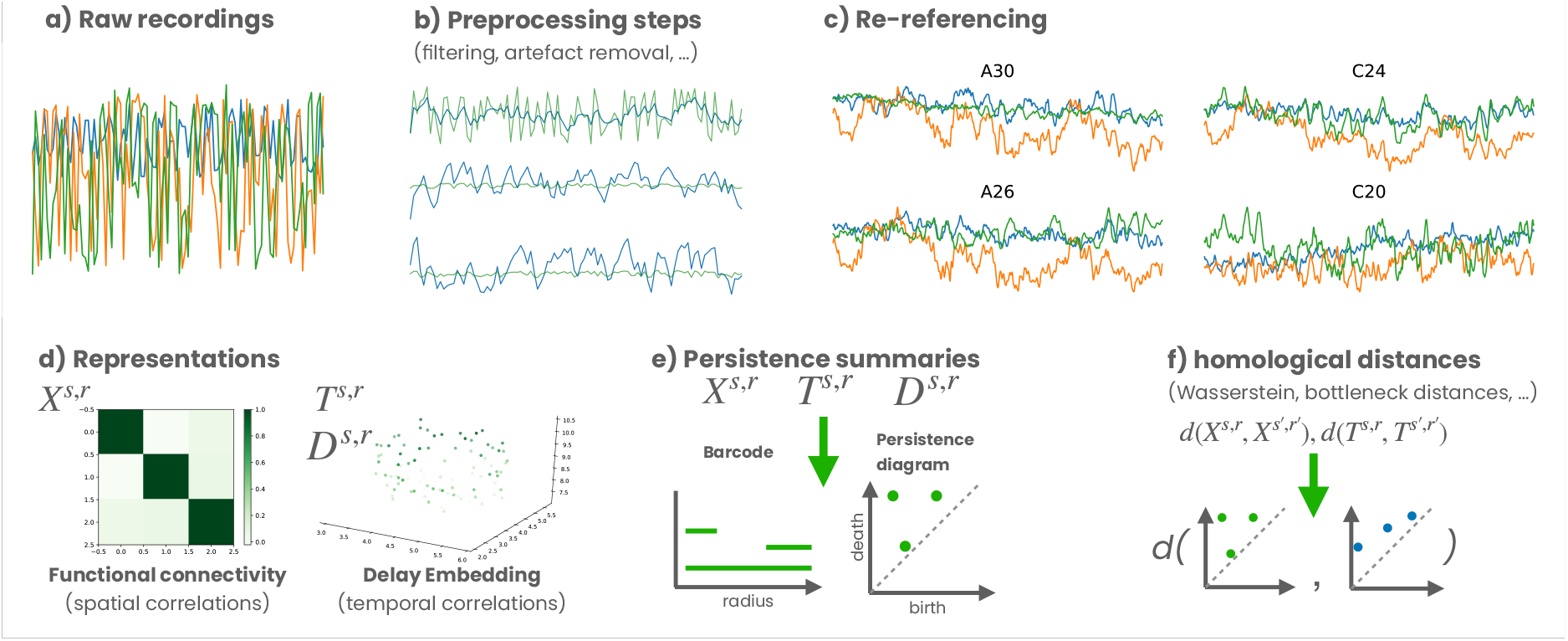
Overview of analysis pipeline. The standard pipeline of EEG analysis follows these steps: a) raw signals are recorded from scalp EEG electrodes. b) the signals are filtered in order to remove noisy or uninteresting frequency bands (here, any activity <0.1Hz and >60Hz, as well as 50Hz electrical line noise) (*filtered*); recorded signals are then cleaned to remove artefacts (i.e. blinks, eye movements, muscle artefacts, heart rate artefacts, electrode pops - i.e. single or multiple sharp waveforms that appear after a sudden change in impedance, electrode drifts due to sweat,etc.) (*clean*); interpolated to account for technical issues (i.e. “dead” electrodes), electrode drifts due to sweat or electrode bridging (when electrolyte gel spreads between adjacent electrodes), etc. (*cleanint*). c) pre-processed data are then referenced to one of the electrodes or, in some case to the average value across all channels. Different reference choices can result in different effects: for illustration, we show here three intervals of EEG signals for four different references; note how the relation among series can change depending on the choice of the reference. d) In this study we investigate three representations: 1) *X* is obtained by considering the Pearson correlations among channels and results in a description of spatial correlations, 2) *T* is the Takens (or delay) embeddings starting from the multivariate EEG timeseries and explicitly encoding temporal correlations within the signals, 3) *D* is a variant of *T* wherein the EEG timeseries is directly embedded, that is, without the imposition of time-delay vectors, effectively equivalent to considering the brain configuration space. e) We analyse the three types of embeddings using persistent homology, which quantitatively captures the shape of generic spaces in the form of barcodes or persistence diagrams. f) Finally, we can associate a distance between spaces by measuring distances between persistence diagrams themselves.

### 2.2 Subjects

We tested twenty-one right-handed participants (18 male, 3 female, age range 21-39, mean age ± standard deviation: 25.76 ± 4.54 years). No participant had a history of or current neurological or psychiatric illness, according to self-report. Data from one participant were excluded due to excessive EEG artifacts, thus leaving 20 participants in the final sample (17 male, 3 female; aged 21-39). All participants provided written, informed consent to procedures approved by the cantonal ethics committee (CER-VD, Switzerland).

### 2.3 Recording Procedure and Pre-processing Pipelines

Participants sat in a sound-attenuated darkened room (WhisperRoom MDL 102126E), and were first tested using a multisensory paradigm. Event-related potentials from this dataset have already been published in (Tivadar et al., 2018). After the experimental paradigm, participants were asked whether they would like a break before the resting state recording was initiated. Participants were then asked to close their eyes and instructed not to engage in any specific physical or mental activity for 3 minutes. Continuous EEG was recorded at 1024Hz with a 128-channel BioSemi ActiveTwo AD-box (www.biosemi.com). No online filters were used. Online references were the typical BioSemi CMS and DRL electrodes, which form a feedback loop that drives the average potential of the subject as close as possible to the amplifier “zero”. Data were offline re-referenced to at least three different references at different pre-processing steps (*filtered, clean*, and *cleanint*, described in more detail later, Figure 1b). We chose those electrodes on the *N* = 128 BioSemi cap that were closest to and most representative of the classical external electrodes used for referencing (Chella et al., 2016). Specifically, we used the average reference as well as C17, A23 and D24B14 as representative of nose, inion, and linked-mastoids/earlobes references, respectively. Prior to cleaning, a 2nd order Butterworth filter (−12dB/octave roll-off; 0.1Hz high-pass; 60Hz low-pass; 50Hz notch) was applied, which was computed linearly in both forward and backward directions to remove phase shifts. Thus, by filtering, any activity lower than 0.1Hz and higher than 60Hz was removed, together with 50Hz activity which is typical of electrical noise (i.e. power line noise). These filtered data (denoted *filtered* in the following) constitute the first pre-processing pipeline we will consider. Next, we further pre-processed the filtered data, by cleaning them (*clean* dataset). Data quality was thus controlled first via visual artifact detection and then via ICA decomposition in Matlab (R2020a) using the EEGLAB toolbox (Delorme and Makeig, 2004), in order to exclude any remaining transient noise, muscle artefacts, heart beat artefacts and lateral eye-movements or blinks. These data were also referenced to three references, excluding the average reference. This constitutes the second pre-processing pipeline. Finally, for the third pipeline (*cleanint*), we further inspected data from artefact-contaminated electrodes. These electrodes were interpolated using spherical splines (Perrin et al., 1989), which take into account all of the electrode sites. We then re-referenced our dataset to all the four references specified above, including the average reference. To summarize, the *filtered* and *clean* data were only referenced to C17, A23, D24B14, while the *cleanint* data was referenced to the previously named electrodes, and additionally to the average reference.

### 2.4 Functional connectivity metric embedding construction

We compute functional connectivity networks using Pearson correlations. More precisely, for each combination of subject *s* and reference *r*, we compute the correlation matrix **C**^*s,r*^, in which entry 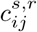 corresponds to the Pearson correlation between the timeseries of channels *i* and *j* with respect to a fixed subject *s* and reference *r*. To each subject-reference pair (*s, r*) we associate to **C**^*s,r*^ a discrete metric space *X*^*s,r*^, obtained by mapping the timeseries corresponding to each channel *i* to a point *p*_*i*_ *∈ X*^*s,r*^. Distances between points are given by 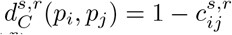. This defines a metric space, 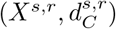) for each subject-reference pair (*s, r*).

### 2.5 Delay embedding construction

Despite the presence of unmodeled noise when re-referencing EEG potentials, the EEG signal samples from the brain’s dynamical state space. To reconstruct that underlying dynamical system, we compute the Takens embedding of each volunteer’s re-referenced multichannel EEG recordings. As before, for each combination of subject *s* and reference *r*, we compute the Takens embedding *T* ^*s,r*^ as follows: for a single time series *x*(*t*) we build a *d*-dimensional point cloud defined as *{x*(*t*_0_), *x*(*t*_0_ + *τ*), *…, x*(*t*_0_ + (*d* − 1)*τ*) *}* for all *t*_0_ in the time series, and where *τ* is a delay and *d* the embedding dimension. Standard techniques are adopted to choose the pair (*τ, d*). Distances between points in Takens embeddings are computed using the canonical Euclidean distance, *d*_*E*_, as prescribed by Takens’ embedding theorem (Noakes, 1991). It is possible to generalize the embedding to the case of *I* time series, which in turn results in a *d × I* dimensional embedding. We therefore consider the metric spaces, 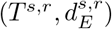). (Myers et al., 2019).

### 2.6 Direct embedding construction

Following Takens’ finding that the delay embedding characterizes a system’s dynamical state space, Deyle and Sugihara (2011) found similar properties when directly embedding the temporal evolution of the *I* dimensional vectors from multi-channel recordings (Deyle and Sugihara, 2011). As before, for each combination of subject *s* and reference *r*, we compute the direct multichannel embedding *D*^*s,r*^ as follows: for a multichannel time series ***x***(*t, i*) we build a *I*-dimensional point cloud defined as {*x*(*t*_0_, *i*_0_), *x*(*t*_0_, *i*_1_), *…, x*(*t*_0_, *i*_*I*_)*}* for all *t*_0_ in the time series, and where *I* is the number of recording channels. Here again, distances between points are computed using the canonical Euclidean distance, 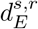. Thus we develop the metric spaces of the direct embedding, 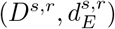).

The benefit of the direct embedding is in its interpretability. Each embedded point corresponds to the vector of signals from all electrodes at a specific time point, an instantaneous EEG activation. However, the number of embedded points increases by a factor of *N/τ* where *N* is the total number of time points in the recording. In order to decrease computation time, and as data were low-pass filtered from 1024 Hz to 60 Hz, data in the multichannel embeddings were downsampled from 1024 samples per second to 64 samples per second.

### 2.7 Topological distances between spaces

We compute distances between spaces using persistent homology (Edelsbrunner and Harer, 2008; Ibáñez-Marcelo et al., 2019b). More precisely, we perform standard persistent homology analysis on the Rips-Vietoris filtrations defined over the points in *X*^*s,r*^, *T* ^*s,r*^, or *D*^*s,r*^. Persistent homology works by studying the evolution of topological features (connected components, 1-dimensional cycles, 3d-cavities, etc.) along a series of progressively finer simplicial complex approximations. A simplicial complex can be intuitively imagined as a higher-dimensional version of a graph, that in addition to edges (that is, pairs of points or vertices, called 1-simplices) also allows for other elementary bricks composed by groups of *k* + 1 points, called *k*-simplices (*k* ≥ 2). In our case, however, we need a way to go from metric spaces to these simplicial complex approximations. We do this using the Rips-Vietoris construction. It works as follows: given a set of points *{p*_0_, *p*_1_, *…, p*_*n*_*}* in a metric space *M* and an arbitrary radius *r*, for each point *p*_*i*_ we consider its neigh-bourhood **B**(*p*_*i*_, *r*) of radius *r*; we define simplices in the Rips-Vietoris complex *RV* (*M, r*) at distance *r* as follows: whenever **B**(*p*_*i*_, *r*) *∩* **B**(*p*_*j*_, *r*) ≠*∅* for some *i, j* we add the 1-simplex [*p*_*i*_, *p*_*j*_]; whenever three points *p*_*i*_, *p*_*j*_, *p*_*k*_ all have non-empty pairwise intersections we add the 2-simplex [*p*_*i*_, *p*_*j*_, *p*_*k*_], and so on for higher dimensions. The collection of all these simplices constitutes *RV* (*M, r*).

The choice of *r* is of course problematic, as it requires picking a scale for the simplicial complex reconstruction. Persistent homology inverts the problem by scanning the properties of *RV* (*M, r*) as a function of *r*. The ordered collection of *{RV* (*M, r*) *}* _*r*_ is called a filtration of *M* (Figure 2a). The outputs of persistent homology are barcodes (and equivalently, persistence diagrams). These compressed summaries recapitulate the homological features of a space, describing how long certain topological features live along the filtration (e.g. connected component, 1-dimensional holes, etc.) (Fig. 2b). Each bar corresponds to a specific topological feature and its birth *r*_*b*_ and death *r*_*d*_ radii correspond to the radii at which that feature first appears and disappears, respectively. Persistence diagrams provide an equivalent description: each topological feature is represented in the 2-dimensional plot by a point with coordinates (*r*_*b*_, *r*_*d*_). We adopt the persistence diagram description, because it makes it easier to compute distances between them, and use those distances as a measure of similarity between the corresponding spaces (Edelsbrunner and Harer, 2008).

**Fig. 2.**
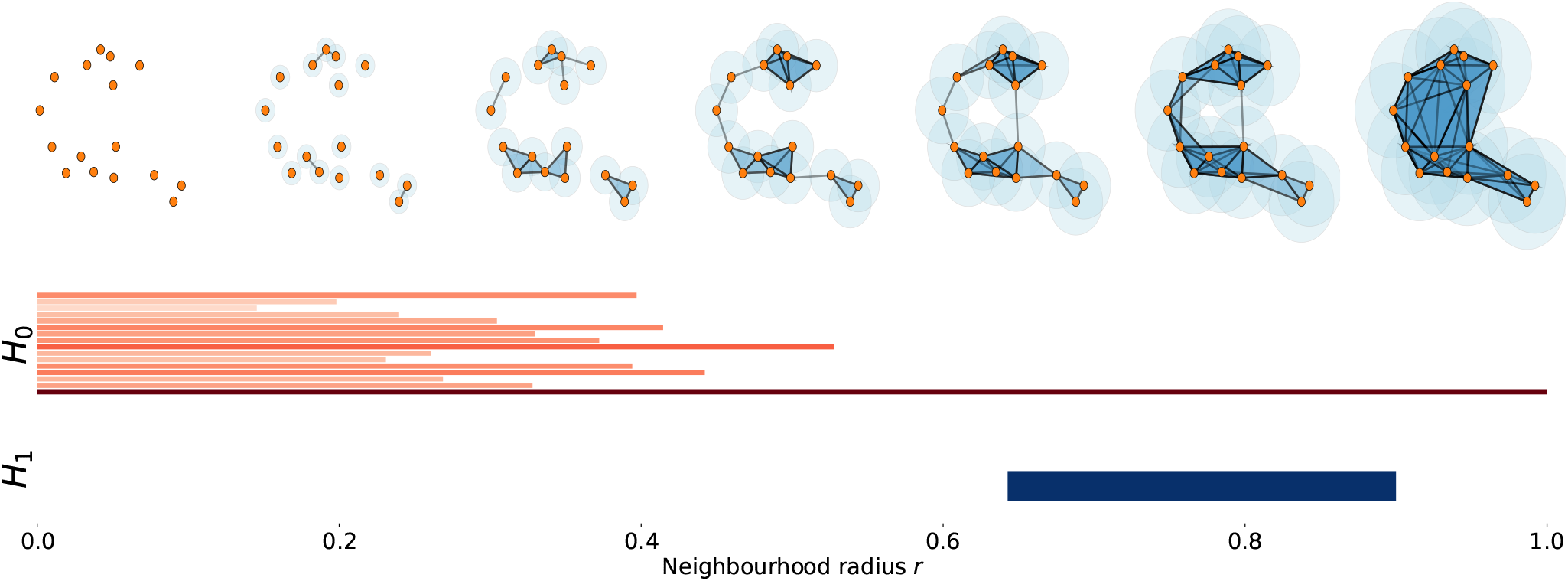
Sketch of persistent homology computation. a) an example of a filtration in two-dimensions. As *r* increases the neighbourhood become larger and more simplices appear, making the Rips-Vietoris complex progressively denser. At the beginning (*r* = 0), the points all belong to components disconnected from each other. As *r* grows, the components begin to merge until only one component remains, which contains all points. Around *r ∼* 0.65, a 1-dimensional cycle appears in the simplicial complex and persists until around *r ∼* 1. b) The barcodes describing the lifetime of the various connected components (red bars), progressively merging into each other until only one survives (describing *H*_0_), and the lifetime of the single 1-dimensional cycle described above (blue bar, describing *H*_1_). Barcodes provide a summary of the topological properties of a space and can be used to compare them in a formal way. We show here the barcode presentece because it makes it easier to relate to the filtration. However, they are equivalent to persistence diagrams, which in turn are more amenable to compute (Wasserstein) distances between spaces.

**Fig. 3.**
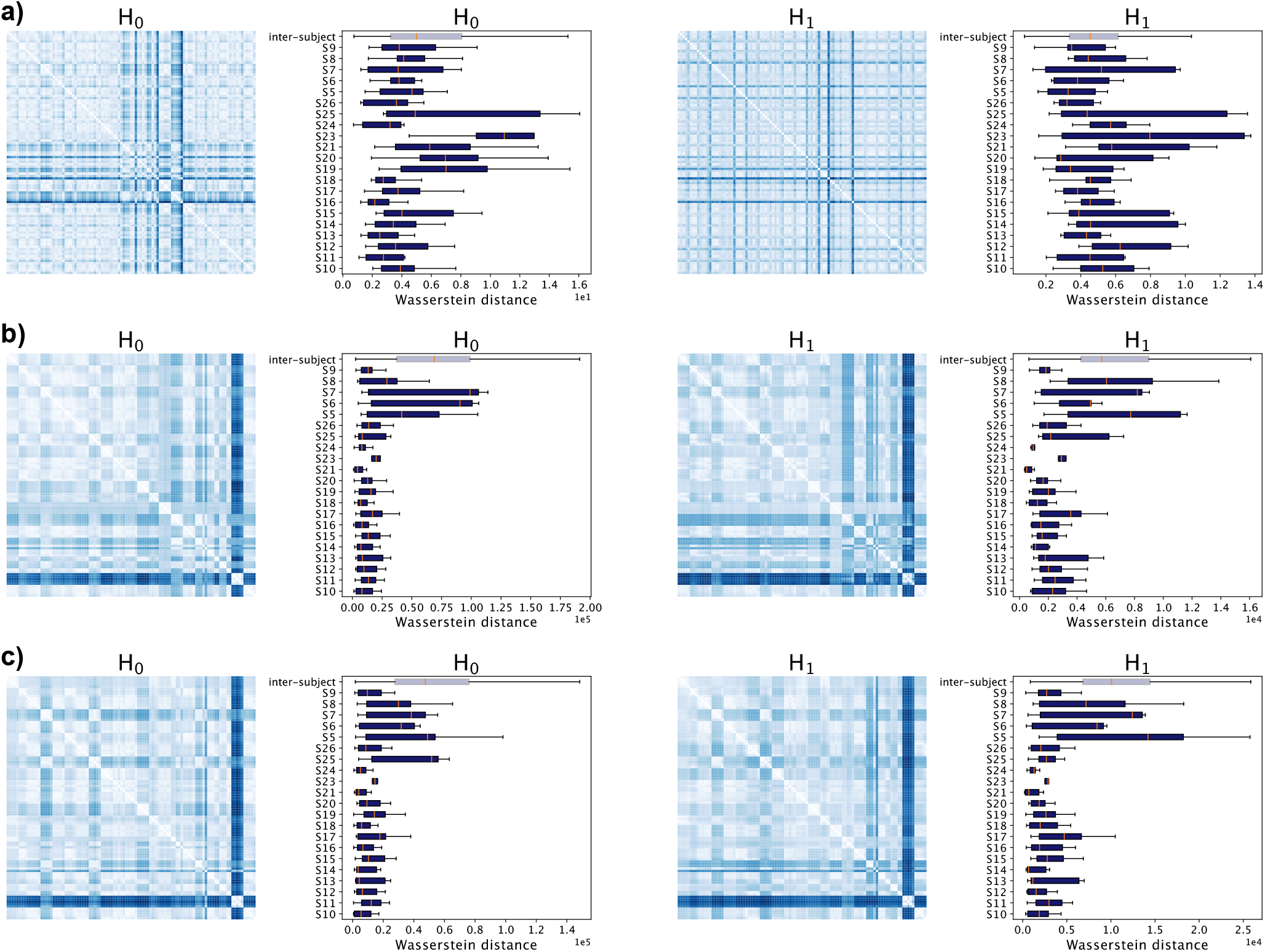
Topological distance between spaces for different references. Panels a) b) and c) refer respectively to the *X*^*s,r*^ *T* ^*s,r*^ and *D*^*s,r*^ embedded spaces for the *cleanint* datasets. For each embedding type, we compute distances between subjects and references for the first two homological groups, *H*_0_ and *H*_1_, using sliced Wasserstein distances between the corresponding persistence diagrams. We show distances between all (*s, r*) pairs in the heatmaps (with distances growing from white to blue). Rows and columns are ordered by subject and then by reference. Therefore, the presence of diagonal blocks of short distances (lighter colors with respect to the off-diagonal blocks) implies that re-referencing induces small changes with respect to inter-subject variability. It is easy to observe and modular block structure for the *T* ^*s,r*^ and *D*^*s,r*^ spaces that is wider than for the *C*^*s,r*^ spaces. Boxplots further support this result: we show the distribution of within-subject distances between spaces corresponding to different references (*d*(*X/T/D*^*s,r*^, *X/T/D*^*s*^*′,r′*)|*s* = *s′∀r, r′*, divided by subject, one dark coloured box for each subject) and compare it to the inter-subject distances (*d*(*X/T/D*^*s,r*^, *X/T/D*^*s*^*′,r′*) |*s* … *s′ ∀r, r′*, lighter color). For the temporal embeddings, the within-subject distances are smaller (KS test, *p <* 0.01) than the between-subject distances. Results for other pipelines are reported in Figures A.1, A.2 and A.3.

In *X*^*s,r*^ spaces we use the Pearson distance as distance between points to construct the filtration(Fulekar, 2009). In *T* ^*s,r*^ and *D*^*s,r*^ spaces we instead adopt the Euclidean distance.

Here, for computational reasons, we focus on the first two homological groups: *H*_0_, that describes connected components, and *H*_1_, that describes one-dimensional holes. Persistence diagrams are equipped with a metric themselves. We can therefore measure distances between them and use this homology-based distance as a topological distance between spaces (Reininghaus et al., 2015). We choose persistent homology as a descriptor for our study because it allows us to compare spaces with different numbers of points, dimensions and metric structure. More precisely, we define the homological distance between **C**^*s,r*^ and **C**^*s*^*′*, ^*r*^*′* to be the sliced Wasserstein distance between the persistence diagrams corresponding to *X*^*s,r*^ and *X*^*s*^*′*, ^*r*^*′*. Similarly, we define the homological distance between Takens embeddings *T* ^*s,r*^ and *T* ^*s*^*′*, ^*r*^*′* to be the Wasserstein distance between the corresponding persistence diagrams. The same applies for computing distances between multichannel embeddings *D*^*s,r*^ and *D*^*s*^*′*, ^*r*^*′*. For completeness, in the following we report the results for *H*_0_ and *H*_1_, although in this study the results from both dimensions typically agree.

### 2.8 Comparison with temporally reshuffled null model

We also consider an auxiliary randomized version of the (*s, r*) datasets: given each dataset

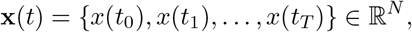

we keep the vectors the same but we reshuffle the temporal labeling. In this way, we preserve the statistical properties of the signal, but destroy the temporal correlations within it. In particular, this type of randomization conserves exactly the network of spatial correlations between electrodes. That is, those captured by the correlation networks **C**^*s,r*^. Similarly, the direct embedding is unaltered by reshuffling the temporal labels. For this reason, we only re-analyse the Takens embeddings constructed from the randomized data, *T* ^*\**^. The utility of the randomised model is that it provides a benchmark for the effect of reshuffling in addition to the observed inter-subject similarity. It also allow us to tease apart the role of the set of realised brain topographies (well described by *D*^*s,r*^) from that of their temporal order.

## 3 Results

### 3.1 Effects of re-referencing on topology of functional representations

We quantified the effects of changing the EEG reference on three different representations: functional connectivity, computed from spatial correlations between electrodes; direct embedding of the brain activations, representing the space of sampled configurations; and the dynamical landscape of brain activity, as reconstructed from temporal embeddings of the signals. More precisely, the first describes how different regions of the brain co-activate and it corresponds to how neuroimaging signals are often studied, both with fMRI (Bassett and Sporns, 2017; Petri et al., 2014) and EEG (Sakkalis, 2011; Ibáñez-Marcelo et al., 2019b). The direct embeddings instead cast EEG signals as points of a high-dimensional static point cloud, which effectively defines the space of realized activations (Donato et al., 2016). Finally, the Takens embeddings are used to reconstruct the structure of the dynamical attractors of dynamical systems and therefore capture the temporal properties of the system (Myers et al., 2019). We analysed data from *n* = 20 subjects, each re-referenced to *R* = 4 different references, using *filtered, cleaned* and *cleaned interpolated* data. For each pair (*s, r*) we computed the corresponding correlation *X*^*s,r*^, Takens *T* ^*s,r*^ and direct *D*^*s,r*^ embedding spaces. We then computed their persistence diagrams as described in Methods and measured the Wasserstein distances between them. In particular, we were interested in quantifying the changes induced by re-referencing data from the same subject. Figure 3a) shows the distances between the *X* spaces computed between all subjects and references, for *H*_0_ (left) and *H*_1_ (right). Figures 3 parts b) and c) show the same for *T* and *D* spaces, respectively.

Data calculated against different references and belonging to the same subject are grouped together in the heatmaps (increasing distances go from white to blue). Thus, diagonal blocks of short distances (see definition below) suggest that re-referencing induces mild changes in the topological structure of the spaces under study. It is important to specify what is meant by short distances. We choose here to use the distance between spaces corresponding to different individuals (that is, *s ≠ s′*) as the benchmark for these effects. This choice is predicated on the finding that the brain waves of each individual are a unique biometric signature for that individual (Poulos et al., 1999; Marcel and R. Millan, 2007; Chan et al., 2018). The boxplots in Figure 3 show this in quantitative form: for each subject (dark coloured bars) we plot the distribution of distances between the persistence diagrams of spaces obtained from the different referencing. In addition, we plot (in lighter color) the distribution of distances between the spaces corresponding to different subjects (for any combination of references *r, r′*).

It is plain to see that the set of intra-subject distances between re-referenced direct and temporal embeddings are generally shorter than distances measured between subjects. By contrast, we see that the set of intra-subject distances between re-referenced correlation spaces *X* are approximately the same value as inter-subject distances. Hypothesis testing using the two-sided Kolmogorov–Smirnov test (*p <* 0.01) confirms that the correlation metric spaces have statistically similar inter-subject embeddings and intra-subject embeddings.

Beyond statistical significance, we can also quantify how dissimilar intra-subject versus inter-subject distances are by computing the effect size between distributions, following Cohen’s *d* method (Kelley and Preacher, 2012). We find that the magnitude of the effect sizes is consistently much larger for the direct embedding spaces *D* and the temporal embeddings space *T* than for the correlation spaces *X* (Figure 4) (Sawilowsky, 2009). Our results therefore imply that the topology of the brain configuration spaces and of the temporal embeddings retain individual-specificity after re-referencing of the EEG data. By contrast, correlation embeddings appear to be non-specific to a given individual subject. Observed differences between temporal embeddings and correlation embeddings are especially apparent in *H*_1_. More generally, these results are consistent with the demonstration of the presence of self-specific temporal states amid a uniform backdrop of group-similar EEG correlation.

**Fig. 4.**
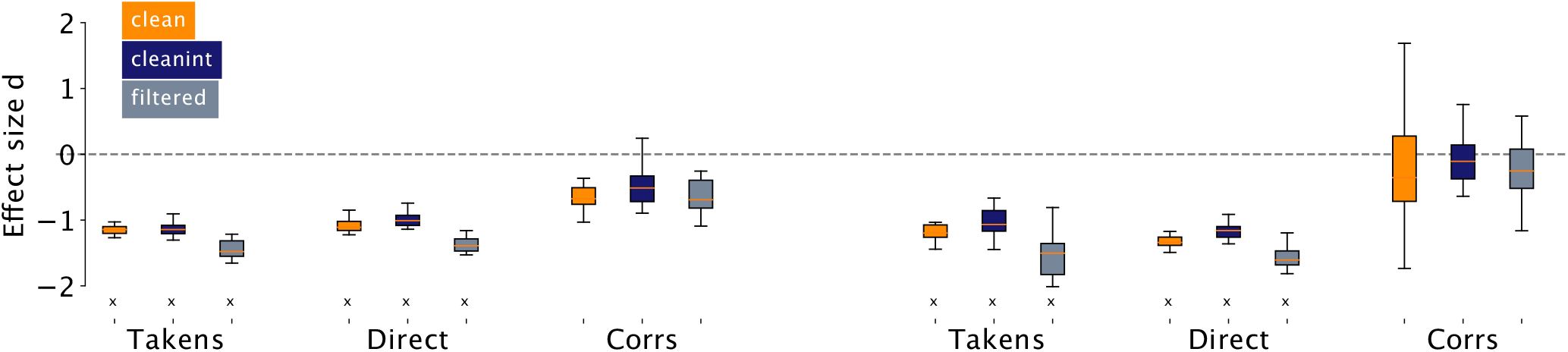
Effect size distributions for within-subject versus inter-subject distances. For each subject *s*, we compute the Cohen’s *d* for the difference between within-subject distances (across references) versus the inter-subject distance distribution. For each pre-processing step *r*, and type of embedding (*T*, T, D, X*), we collect the Cohen’s *d* values across all subjects and display them as a distribution. Instances where the set of within-subject distances are significantly (*p <* 0.05) different from inter-subject distances are marked with an ‘x’. Instances where the distribution of real Takens embeddings *T* ^*r*^ are significantly different from the distribution of randomized Takens embeddings *T\** ^*r*^ are marked with a *†*. The effect of pre-processing is to remove idiosyncratic outliers. We find that for all studied pipelines, the Takens embeddings *T* ^*s,r*^ show larger differences (larger absolute effect size) with respect to the *X*^*s,r*^ spaces, implying that Takens embeddings are more robust to re-referencing than functional connectivity.

### 3.2 Role of different pre-processing pipelines

We next investigated whether the above results vary significantly for different choices of pre-processing. Indeed, EEG signals usually undergo a series of steps before being considered suitable for analysis. We repeated the analysis for all the pipelines.

Similarly to what we did in the previous section, we show the results for effect sizes in Figure 4. We find that effect sizes are always larger (in magnitude) for the *T* ^*s,r*^ and *D*^*s,r*^ spaces than for the corresponding *X*^*s,r*^ spaces, and do not differ much across pre-processing pipelines (*d ∼* 1 − 1.5, considered a very large effect (Sawilowsky, 2009)). This result suggests that different pre-processing pipelines do induce some changes in the reconstructed topology, but also that these changes are small compared to what happens in the case of correlation networks. Additionally, we confirm the result that re-referencing induces changes in the reconstructed topology of *D* and *T* spaces that are much smaller than the inter-subject variability.

### 3.3 Comparison with the temporally reshuffled null model

We showed that *T* ^*s,r*^ and *D*^*s,r*^ spaces display stronger topological robustness with respect to those built from spatial correlations, while at the same time appearing to be more subject-specific. However, we have not ascertained whether these properties are a consequence of the statistics of the signals themselves, or rather they truly emerge from their temporal features. We tested these alternatives by comparing previous results to a null model in which we destroy temporal correlations by reshuffling the time label of instantaneous activity. Note that –by construction– the matrices **C**^*s,r*^ (and thus the *X*^*s,r*^ spaces) remain exactly the same under this reshuffling. In other words, this means that we preserve exactly the spatial correlations, while we destroy the temporal ones. The same argument holds for the *D*^*s,r*^ embeddings: it is a direct embedding from the multi-channel EEG signals and therefore reshuffling the temporal labels does not change the point cloud. Here, we therefore only focus on the effects on *T* ^*s,r*^ spaces, by investigating the changes induced in the Takens embeddings constructed from the reshuffled series, 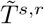 spaces.

In Figure 5a) and b) we compare the results for *H*_0_ and *H*_1_. The first observation is that the distances between reshuffled datasets corresponding to the same subject (across different references) appear to be much more heterogeneous than in the case of real data. This can be observed in several different ways. The distances between references of the same subject are much farther away from each other in the reshuffled case than in the real case (solid color boxes versus the corresponding light colored boxes). Similarly, the inter-subject distances are typically larger in the reshuffled cases. This also holds for the distances between a specific real (*s, r*) pair and its reshuffled version (white box, labeled as *real-rand* in Fig. 5a-b). In fact, this latter distance distribution (*real-rand*) and the reshuffled inter-subject distance distribution (for all pipelines) have averages that are statistically indistinguishable (Mann-Whitney *u* test for equal mean, null hypothesis not rejected at *p <* 0.01), while the real inter-subject distance has a smaller average (significant on the same test). The same results hold for the other pre-processing schemes (Figure A.4).

**Fig. 5.**
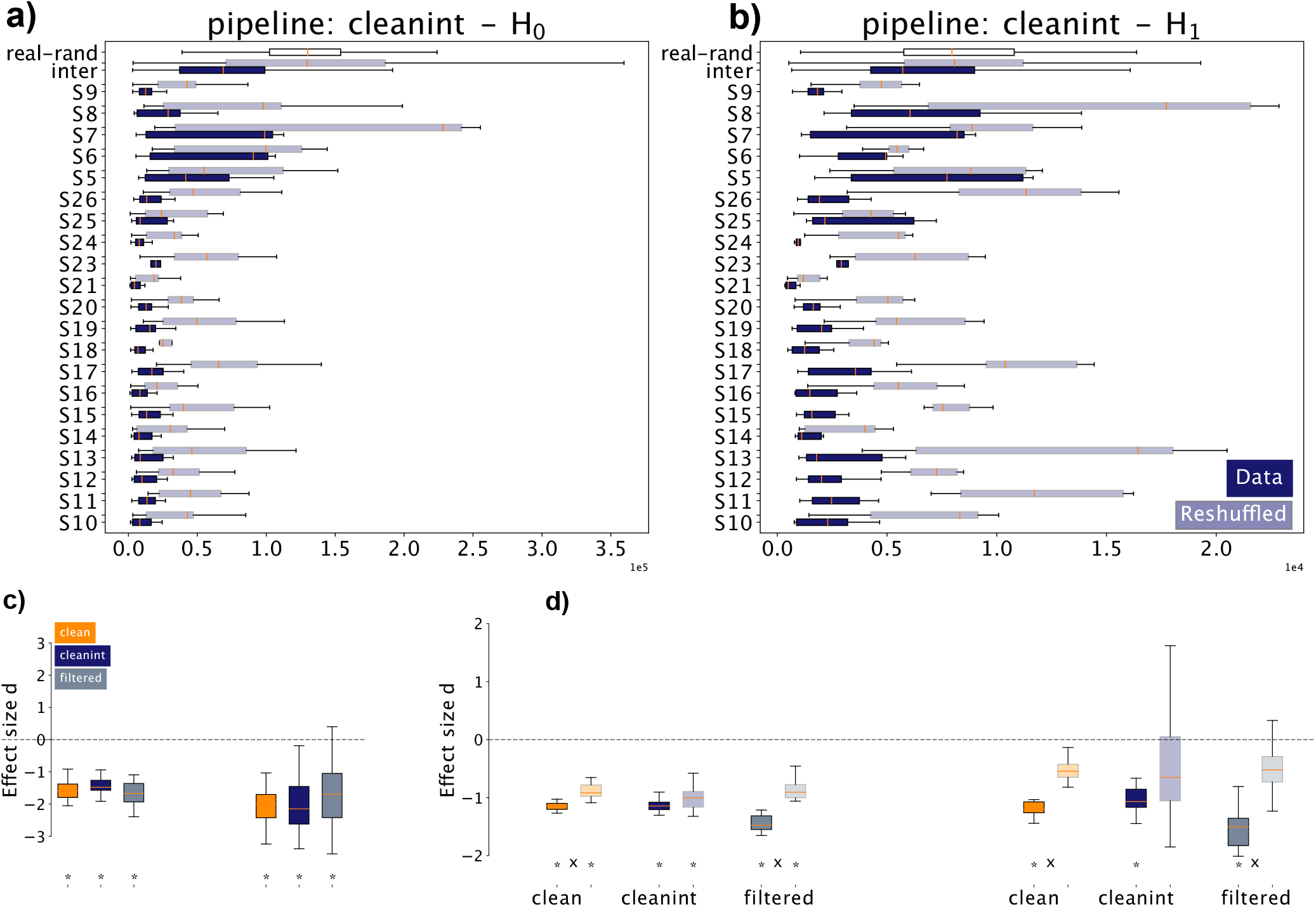
Comparison of real with reshuffled data. *a*) and *b*) Distance distributions (respectively for *H*_0_ and *H*_1_) between references of individual subject (solid color), between the temporally reshuffled data (lighter color boxes). *inter* labels the inter-subject distance distribution for both real (solid) and reshuffled data (lighter color). *real-rand* (white box) represents the distribution of distances between a (*s, r*) pair and its temporally reshuffled version. Distances among references of the same subject have generally smaller mean and variance with respect to the reshuffled data. Moreover, the distances between a (*s, r*) pair and its randomized versions are often larger than those between different subjects. *c*) we confirm this by computing the corresponding effect size via Cohen’s *d*, that is the effect sizes of the distances *d*(*T* ^*s,r*^, *T* ^*s,r*^*′*) versus that of *d*(*T* ^*s,r*^, 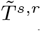). For both *H*_0_ and *H*_1_ and for all pipelines, the effect sizes across subjects are significantly smaller than zero (varying between −1 and −2, considered to be very large effects, asterisks indicate significance on one-sample t-test at *p <* 0.01 Bonferroni corrected for multiple comparisons to reject the null hypothesis that the effect size mean is 0). *d*) for all pipelines, effect sizes for the intrasubject distance distributions *d*(*T* ^*s,r*^, *T* ^*s,r*^*′*) versus the inter-subject *d*(*T* ^*s,r*^, *T* ^*s*^*′,r′*). Solid color boxes indicate comparison between real data, lighter color boxes indicate the same comparison for the reshuffled data. Asterisks indicate significance on one-sample t-test at *p <* 0.01 Bonferroni corrected for multiple comparisons to reject the null hypothesis that the effect size mean is 0; crosses indicate indicate significance on Mann-Whitney *u* test at *p <* 0.01 Bonferroni corrected for multiple comparisons to reject the null hypothesis that the real and reshuffled samples have the same mean. In most cases, the effect sizes are significantly smaller than 0, confirming that the real distance distributions have smaller mean with respect to the inter-subject distances with respect to the corresponding reshuffled data.

We further support these observations by computing the corresponding effect size via Cohen’s *d*, that is the effect sizes of difference between the distributions of distances *d*(*T* ^*s,r*^, *T* ^*s,r*^*′*) versus that of *d*(*T* ^*s,r*^, 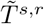). For both *H*_0_ and *H*_1_ and for all pipelines, the effect sizes across subjects are significantly smaller than zero (Figure 5c). Hence, destroying temporal correlations induces varied and heterogeneous changes in the topology of the resulting embeddings; changes which are much larger than those induced by pure re-referencing.

Finally, we ask how large are the changes induced by re-referencing with respect to intersubject variability. Similarly to Figure 4, we compute the effects size for the distribution of intra-subject and inter-subject distances for the real and reshuffled data (Figure 5d).

That is, for *H*_0_ all pipelines display effect sizes significantly different from zero for both real and reshuffled data (one-sample t-test for mean equal 0, *p <* 0.01). For *H*_1_, only real data show an effect size significantly smaller than 0 (same test). Interestingly, when directly comparing the real and reshuffled effect sizes, we find significant differences (Mann-Whitney *u* test at *p <* 0.01 Bonferroni-corrected) only for the *cleanint* and *filtered* datasets. Note that this does not mean that in the *cleanint* datasets the actual distances are the same. Rather, the effect size between the intra-subject and inter-subject distribution is similar in the real and reshuffled cases in the presence of further pre-processing.

Overall, this pattern suggests that the differences previously observed before originate partially from the statistical properties of the signals, irrespective of the time ordering, more precisely in what concerns the relative difference across subjects with respect to different references of the same subject. However, temporal correlations are crucial to constrain the set of possible topologies describing a subject and to discriminate between subjects even when the statistical properties of the signals are similar.

## 4 Discussion

We studied the topological structure of different representations of resting state EEG signals with a particular focus on how the choice of the reference alters the resulting topological observables. We examined three representations that capture different features of the data. The first one was the functional connectivity between electrodes, which we computed as Pearson correlations between electrode timeseries. When computing these correlations, time is integrated away and for this reason the resulting correlations capture the patterns of spatial coactivations among signals at different electrodes. The second type of representation was direct embeddings. In these embeddings, each time point is associated with the vector of instantaneous EEG signal. Together, all the vectors of instantaneous EEG activations define a space that captures the possible configurations of brain activations. Note that in this construction, relative temporal information is lost. Thus, the direct embedding does not encode brain dynamics, but rather the range of possible topographies. Finally, the third type of representation was given by Takens embeddings. They are constructed by concatenating instantaneous EEG vectors corresponding to successive time points. In this way, they reconstruct the properties of the attractor space of a dynamical system (Noakes, 1991; Myers et al., 2019).

We found that the extent of topological changes across correlation spaces corresponding to different references are often comparable with those measured between different subjects. We found that the direct embeddings and Takens embeddings exhibited limited changes across different references and were able to discriminate better across subjects.

The implications of these results are multi-fold. As mentioned above, for a fixed reference, the *D*^*s,r*^ space is the configuration space (i.e. phase space) of EEG whole-brain activations (i.e. topographies (Tivadar et al., 2019)). The facts that the topology of signals does not change significantly across references and that it is subject-specific, suggest that the overall shape of EEG configuration space holds information about the specificity of the underlying individual’s brain dynamics, similarly to what has been observed for simpler dynamical systems (Donato et al., 2016).

However, by construction, the configuration spaces above neglect the role of time, i.e. the temporal ordering in which brain activity appears. It is reasonable then to ask whether the specific order of time points plays a role. The Takens embeddings explore exactly this question. In fact, as mentioned, they allow us to probe the brain’s dynamical attractor space underlying the observed activations (Myers et al., 2019). If there were no information in the temporal ordering, the topological structures of *T* ^*s,r*^ and of its randomized 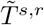 spaces should be similar to one another. Instead, we observed a large standard deviation within embeddings when temporal correlations were removed. More precisely, we observed that a much larger and more heterogeneous set of topological spaces is explored when only spatial correlations and statistical properties of the signals are preserved. Conversely, this implies that temporal correlations constrain to a large degree the set of possible dynamics. These in turn shape how the space of activations differs across subjects’ resting state activity.

There are several ways by which to intuitively understand these findings. One can think the configuration space of activations *D*^*s,r*^ as a mountainous landscape, where each point represents a state that the brain has accessed during the recordings. The Takens embedding instead describes how an individual explores this configuration landscape, i.e. the order of states as explored by a subject; each point in the Takens embedding can be thought as composed by a set of successive positions in the configuration space (a path in the mountainous landscape across different states), that is, a trajectory through time. We found here that both the landscape and how it is explored –the set of trajectories– change across subjects. However, both of these features are very robust to re-referencing and pre-processing choices.

We claim that our results extend to the configuration spaces and the set of trajectories of topographies. While we can only access information about topographies via their referenced instances, our results imply that different reference choices are essentially equivalent from a topological perspective. That is, for different references we find very similar topologies, which therefore implies that we are capturing the topology of the actual space of topographies. We propose therefore to augment conventional topographic analyses with an additional –topological– level of analysis that links together the two levels of description, rather than focusing on a single one at a time.

Naturally, our work also leads to many novel questions, both technical and theoretical: do the topographical configuration space and/or its dynamical properties change under different conditions, e.g. wake versus sleep, altered states, performance of different tasks, etc.? Do shared topological structures emerge under such conditions that are stronger than inter-individiual variability? Previous findings with fRMI and EEG suggest that during tasks the spatial correlation structure is already sufficient to discriminate between tasks, subjects (Ibáñez-Marcelo et al., 2019b,a) or altered states (Petri et al., 2014). It would be indeed important to ascertain which features of the topographic and topological spaces are preserved under different conditions, both analytically and empirically Haufe and Ewald (2019), as this would have direct impact on brain fingerprinting and on functional neuro-degeneration tracking among others (Bari et al., 2019; Rajapandian et al., 2020). This is an exciting endeavour that is currently at the forefront of our current ongoing investigations.

## Authors contribution

M.M.M., B.F. and G.P. conceived the study. R.T. gathered and pre-processed the data. J.B. structured and carried out the topological data analysis over the three embedding types. B.F., G.P.,J.B. and R.T. interpreted together the results. R.T. and J.B. wrote the first draft of the manuscript. B.F. supervised the neuroscientific contents and G.P. the topological ones. All authors contributed to the final draft and review.

## A Appendix

### Results for additional preprocessing pipelines

**Fig. A.1.**
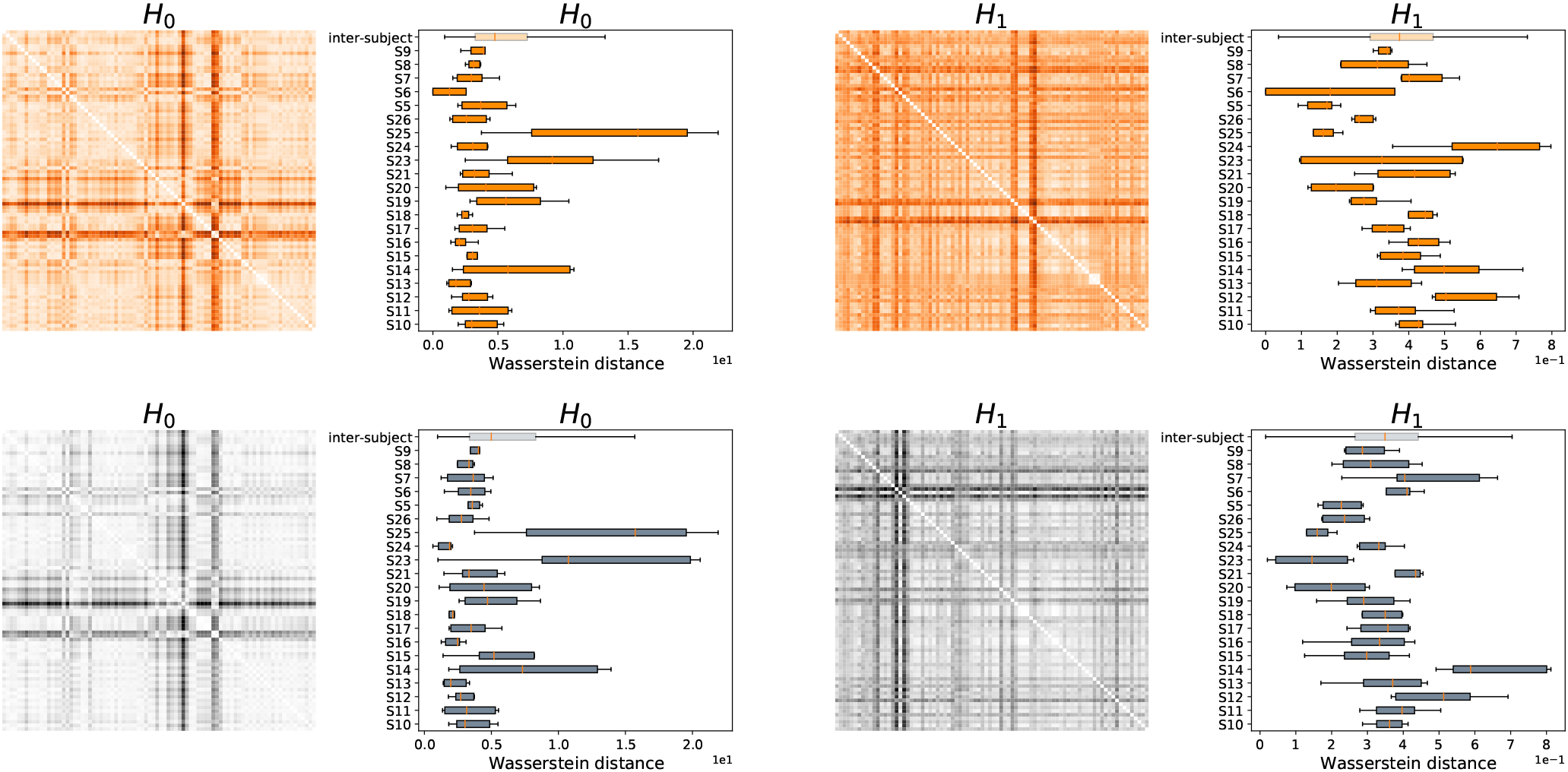
Effect of different preprocessing pipelines on correlation spaces *X*^s,r^. Additional pipeline results for Figure 3: (top row) *clean* pipeline. (bottom row) *filtered* pipeline.

**Fig. A.2.**
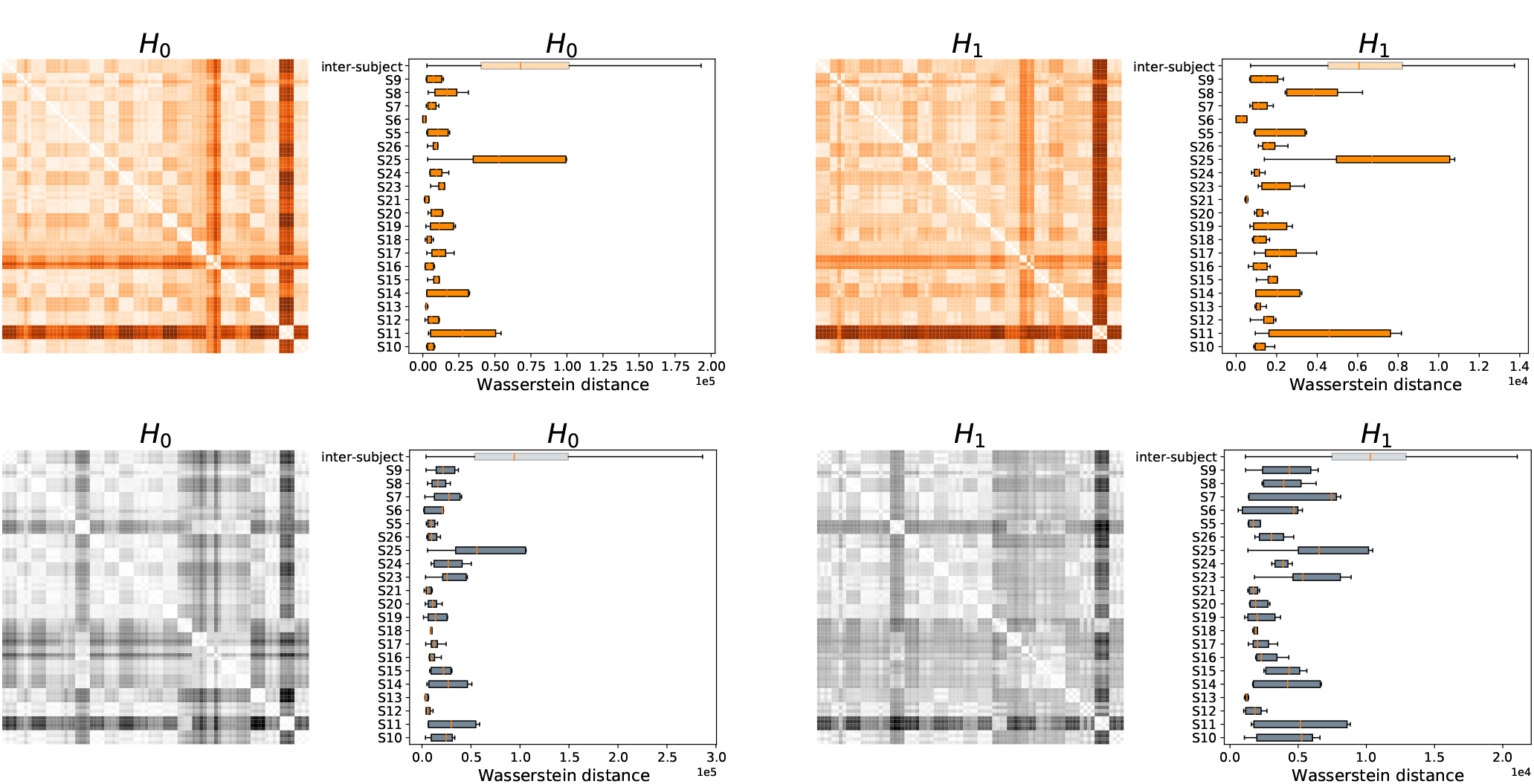
Effect of different preprocessing pipelines on Takens embedding spaces *T* ^*s,r*^. Additional pipeline results for Figure 3: (top row) *clean* pipeline. (bottom row) *filtered* pipeline.

**Fig. A.3.**
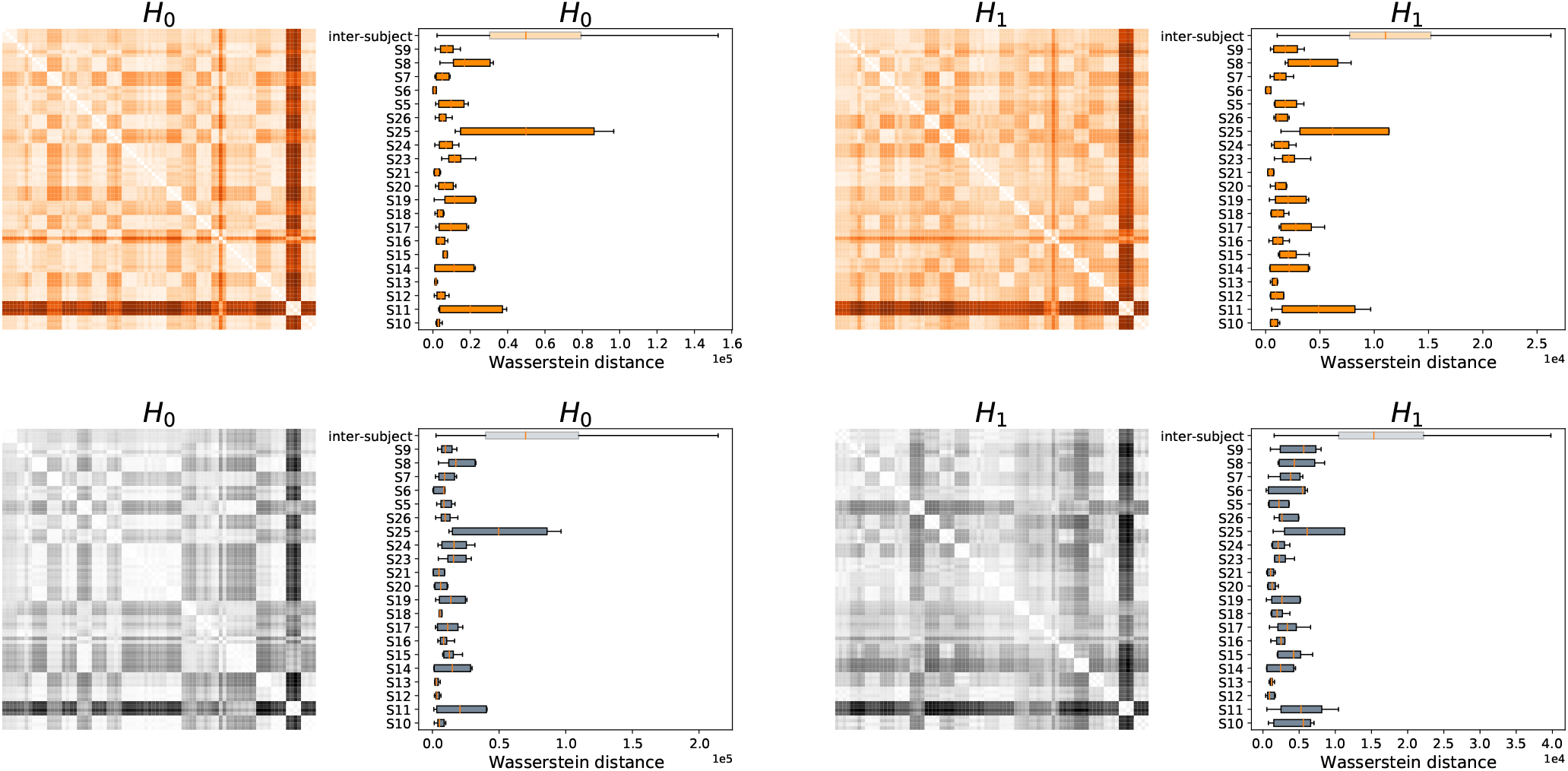
Effect of different preprocessing pipelines on direct temporal embedding spaces *D*^*s,r*^. Additional pipeline results for Figure 3: (top row) *clean* pipeline. (bottom row) *filtered* pipeline.

Results for *T* ^*s,r*^ spaces from temporally shuffled timeseries

**Fig. A.4.**
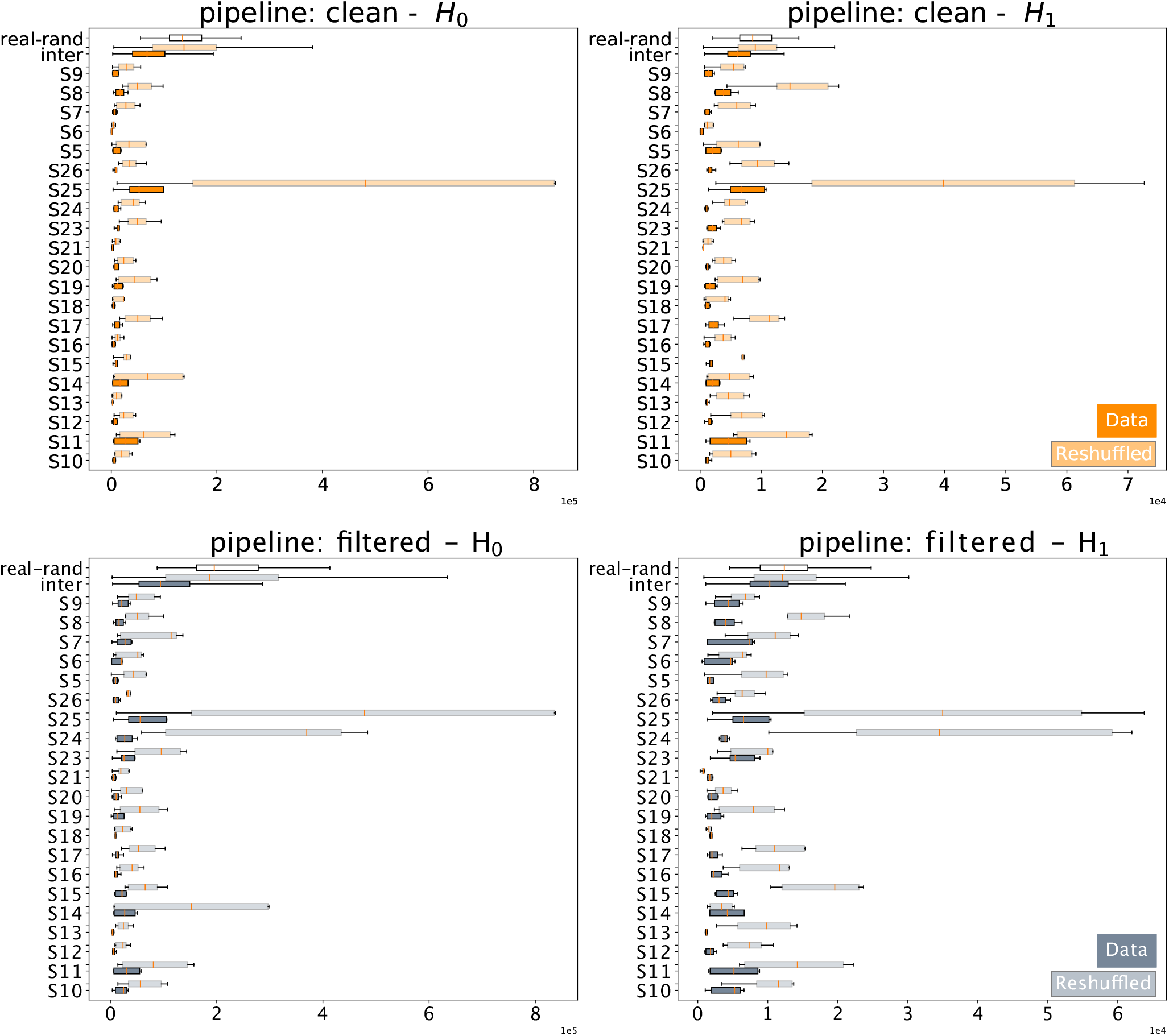
Effect for temporally reshuffled timeseries for *clean* and *filtered* pipelines.

